# Engineering a probiotic *Bacillus subtilis* for acetaldehyde removal: A *hag* locus integration to robustly express acetaldehyde dehydrogenase

**DOI:** 10.1101/2024.06.25.600659

**Authors:** Chandler Hassan-Casarez, Valerie Ryan, Bentley Shuster, John W. K. Oliver, Zachary D. Abbott

**Affiliations:** ZBiotics Company, 44 Montgomery St. 3rd Floor, San Francisco, CA 94104

## Abstract

We have identified solutions to key challenges in probiotic design to create a commercially viable strain for the removal of the intestinal toxin acetaldehyde. Here we report engineering of a σ^D^-dependent flagellin expression locus, hag, as a stable location for robust enzyme production. We demonstrate constitutive gene expression in relevant conditions driven by the endogenous hag promoter following a deletion of the gene encoding a post-translational regulator of σ^D^, FlgM, and a point mutation to abrogate the binding of the translational inhibitor CsrA. Reporter constructs demonstrate hag locus activity after germination and show a steady increase in heterologous expression throughout outgrowth and resumption of vegetative growth. To test the chassis as a spore-based probiotic solution we identified the physiologically relevant pathway of ethanol metabolism and the buildup of gut-derived acetaldehyde after alcohol consumption. Herein, we describe the integration of a robustly expressed Cupriavidus necator aldehyde dehydrogenase, AcoD, under a flagellin protein promoter at the hag locus and report the rapid reduction of acetaldehyde levels after germination in gut simulated conditions.

## Introduction

How we live and experience life including our sleep [1], weight gain [2,3], mental health [4,5], immunity [6,7], and aging [8,9] is linked to the intestinal tract and the gut microbiome. The mechanisms of these relationships involve a complex interplay between proteins and small molecules from our body and our microbiome [10,11]. The intestinal tract is easy to add to (food and small molecule drugs) but challenging to take away from *in situ*. Nature can clear active small molecules in the intestine by the production of enzymes to catalyze conversion. However, the human body and microbiome can lack the necessary enzymes to convert molecular byproducts, especially in the context of modern life, such as from processed foods, environmental pollution, reduced sleep, and increased daily stress. Purified enzymes are a potential solution. However producing and purifying enzymes can be expensive, and pure enzymes are often sensitive to hostile digestive conditions *in vivo* and/or require a cofactor for function. Engineered probiotics are a desirable solution to these limitations, enabling *in situ* regeneration of cofactors, regeneration of proteins, protection from proteases inside the bacterial cell, and low cost of manufacturing that minimizes downstream processing associated with protein purification.

We propose here that successful delivery of active enzymes from engineered probiotics can be achieved with a transient overlay of a non-commensal, endospore-forming, food-safe strain of *Bacillus subtilis*. *B. subtilis*, a Gram-positive, rod-shaped bacterial species, is recognized for its safety and ability to form robust endospores (referred to herein as “spores”), facilitating its survival in harsh environmental conditions [12,13]. As soil microbes, species of *Bacillus* have been ingested by animals for millions of years, are often consumed in human food, and are commonly isolated from human samples [14,15]. In recent years, recognition of the *Bacillus* genus as a probiotic has expanded with *Bacillus coagulans* and *B. subtilis* strains such as HU58 and DE111 (now assigned as *Bacillus inaquosorum*[16]) becoming widespread in foods and supplements as added ingredients [17–19].

The spore-forming nature of *B. subtilis* not only underlines its survivability but also its utility as a probiotic where it can traverse the hostile acidic environment of the stomach. As a matter of product logistics, the stability of bacterial spores also facilitates room temperature storage and a long shelf life. In the vegetative state, *B. subtilis’s* well-characterized nature exhibits strong expression loci, low mutation rates, and high capacity for enzyme production, rendering it an attractive candidate for synthetic biology [20]. With its long history of safe use as a probiotic and its genetic tractability, *B. subtilis* is an ideal platform for engineered probiotics for human consumption.

For probiotic design, to increase safety and stability, it is desirable to minimize edits and exogenous regulatory DNA. Current engineering tools for *B. subtilis* include promoters, secretion tags, and methods for scarless transformation using natural competence [21]. We sought to expand the genetic toolkit in Bacillus by providing a native promoter locus capable of stable high expression, while only requiring exchange of the gene of interest. We’ve identified the *hag* locus, traditionally known for housing the gene encoding flagellin, as a novel site for genomic integration. Supported by initial work from Dr. Dan Kearn’s lab showing a similar approach, we identified the flagellar operon as a candidate locus that highly expresses a non-essential protein during the motile metabolic state of vegetative cells while retaining genomic stability [22,23]. The *hag* gene has a robust endogenous transcriptional promoter and a ribosome binding site that evolved to produce hundreds of thousands of flagellin proteins in a single bacterium [24,25]. *B. subtilis* regulates motility by a sophisticated system involving several positive and negative regulators in a manner that can switch on and off depending on nutrient availability [26]. In this work we share a design where minimal engineering enables robust constitutive expression in tested conditions from the *hag* locus by removing two known repressors. In brief, transcription is mediated by the alternative sigma factor σ^D^, which is bound to and inhibited post-translationally by the FlgM protein [27]. Deletion of *flgM* increases activity of σ^D^, and results in higher and more constitutive transcription of the flagellar operon and specifically the *hag* gene. Additionally, translation of the *hag* gene is enabled by a ribosome binding site that can be bound and repressed post-transcriptionally by a protein called CsrA [28]. A single point mutation in the CsrA binding site abrogates its binding and results in more robust translation of protein from that locus [29].

We leveraged the endogenous promoter of the *hag* locus with a point mutation in the CsrA binding site to generate P*_G->A_* (henceforth referred to as P*_hag*_*), and we deleted *flgM* to improve transcription through this promoter. As a model application for this expression system *in vivo,* we identified acetaldehyde (AcA) in the gut as a target of considerable public interest. After consumption of alcohol, the major pathway for alcohol removal from the bloodstream is catalyzed in the liver by alcohol dehydrogenase (ADH) oxidation of ethanol into AcA. AcA is subsequently oxidized in the liver to acetate via AcA dehydrogenase (ALDH). When ethanol is consumed at a rate that exceeds clearance by the liver it recirculates leading to increased blood alcohol content. Alcohol can equilibrate to the intestine from blood alcohol content or traverse to the colon with food prior to being absorbed into the bloodstream in the first place. When this occurs, intestinal flora can convert colonic alcohol into AcA with microbially expressed ADH, but they do not as efficiently convert AcA into acetate. Gut-derived AcA has been shown to increase in a dose dependent manner *in vitro* and is released back into the bloodstream [30]. In addition, during alcohol ingestion, the highest levels of AcA in the body are found in the colon *in vivo* [31,32]. AcA is well-documented as causing much of the common next-day discomfort associated with alcohol consumption, and its removal has been demonstrated to reduce this accompanying discomfort [33]. Conversely, when the body’s ability to oxidize AcA to acetate is inhibited either chemically (e.g. with disulfiram) or genetically (e.g., single nucleotide polymorphisms in AcA dehydrogenase genes), amplified discomfort is typically experienced [34,35].

In this study, we present a strain with a genomic integration at the *hag* locus, capitalizing on an engineered version of its endogenous promoter to drive robust expression of ALDH enzymes and promote metabolism of gut-derived AcA (Fig 1). We heterologously expressed *acoD* (Uniprot:<u>P46368</u>), a gene native to *Cupriavidus necator* encoding an enzyme with ALDH activity [36]. A successful demonstration of the *hag* locus as an effective site for genomic integration broadens the *B. subtilis* genetic toolkit and accentuates its potential to address real-world challenges through synthetic biology applications. In addition, by evaluating the heterologous expression after germination and in simulated intestinal media, we can extrapolate the practical utility of this engineered chassis as an engineered *in vivo* protein delivery system. We aim for our findings to serve as a steppingstone towards establishing the *hag* locus as a reliable and versatile site for gene integration, paving the way for more functional exploration in *B. subtilis*.

**Fig 1.**
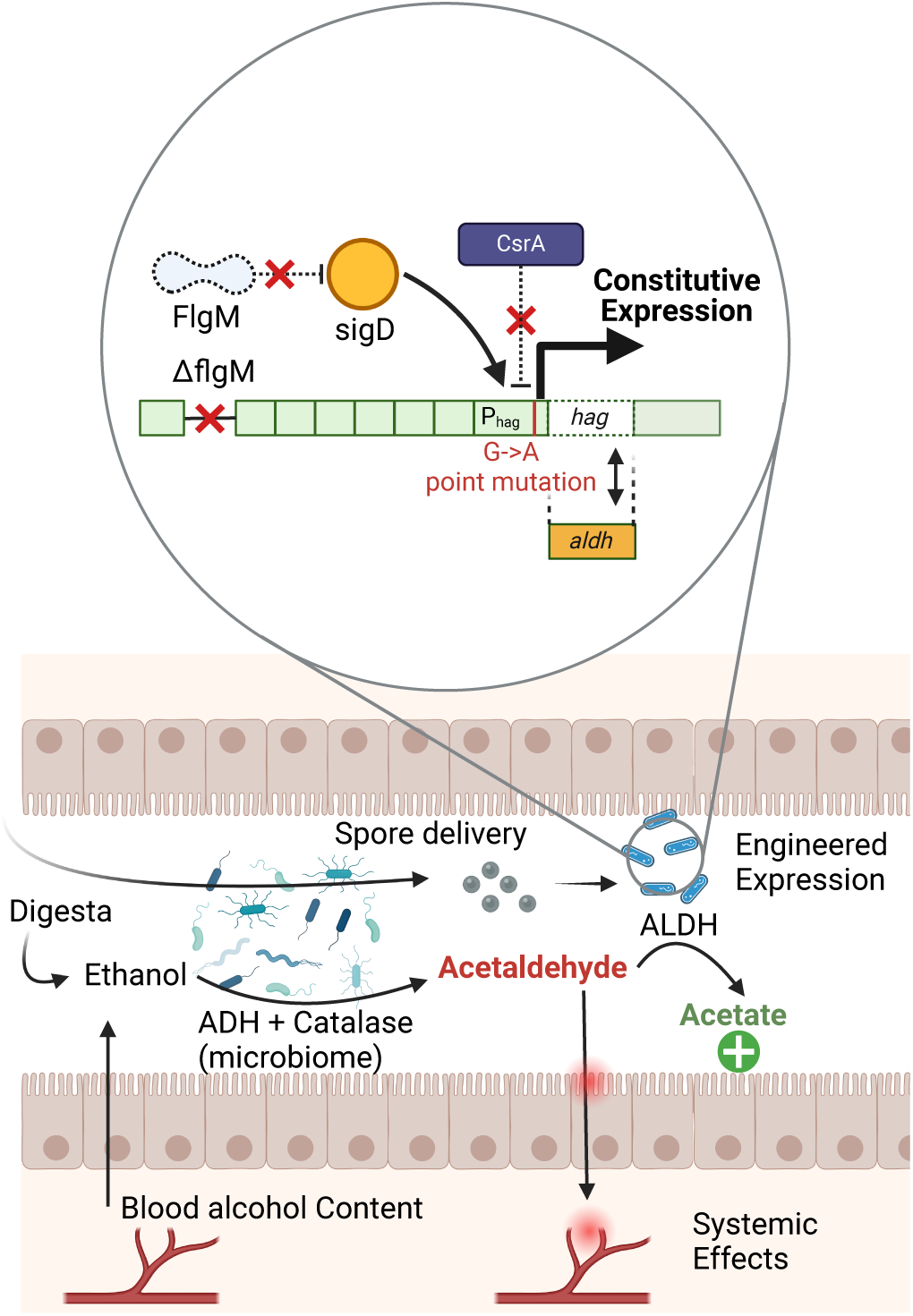
An overview of the engineered promoter and ALDH action in the gut. **Top:** Optimization of *hag* locus engineering for constitutive and robust heterologous protein expression. Deletion of *flgM* alleviates repression of σ^D^ for constitutive transcription, and a point mutation in the *hag* promoter prevents CsrA from repressing translation at the *hag* locus. Heterologous expression of ALDH at this site allows the bacteria to catalyze the enzymatic oxidation of acetaldehyde (AcA) into acetate. **Bottom:** Translational application of engineered chassis for AcA to acetate conversion in the intestine. A small percentage of consumed ethanol can be converted into AcA by ADH-expressing microbes in the gut microbiome prior to absorption and processing in the liver. The AcA can build up in the lumen of the gut and subsequently spread throughout the body. Consumption of engineered *Bacillus subtilis* endospores leads to germination and constitutive expression of ALDH, facilitating the conversion of AcA to acetate before buildup and dispersal can occur.

## Results

### The hag promoter enables constitutive and robust protein expression

In *B. subtilis*, the *hag* locus is an attractive site for genomic integration where it exhibits robust expression of nonessential proteins. As a demonstration of its utility, we aimed to engineer a strain of *B. subtilis* to express acetaldehyde dehydrogenase (ALDH) that is capable of metabolizing gut-derived acetaldehyde. To be effective for this use case, expression would ideally be constitutive so that expression would not be dependent on growth-phase or conditions that may be difficult to predict in the intestines of a diverse population of people. To this end, we modified the promoter of the *hag* locus to relieve repression due to FlgM and CsrA to generate P*_hag_*_*_.

To rapidly assess the overall efficiency of the derepressed engineered P*_hag*_* promoter at the *hag* locus, expression of GFP at that locus was compared to expression at the *amyE* locus using three common robust heterologous promoters: P*_veg_*, P_lac_, and P*_43_* [37–39]. A qualitative comparison of green fluorescence on plates showed that expression through the P*_hag*_* promoter in the *hag* locus greatly exceeded expression at the *amyE* locus in our hands (data not shown), demonstrating the potential for heterologous expression of our engineered strain. To further assess expression from the *hag* locus, a LacZ reporter strain, ZS456 (Δ*flgM* P*_hag*_* Δ*hag*::*lacZ*), was constructed to enable high sensitivity colorimetric measurements. We quantified LacZ activity from germinating cultures. Spores of this strain completed germination by T_30_, as observed by a drop in absorbance at 600 nm (optical density, OD_600_) (Fig 2). OD_600_ increased thereafter as the cells entered the outgrowth phase of spore revival (Fig 2B). LacZ activity was first detectable at T_45_ matching the expected expression timeline delay of protein synthesis for non-essential systems during and after germination (Fig 2A) [40]. Activity increased linearly thereafter up to T_90_, after which we observed a sharp rise as outgrowing cells prepared for vegetative growth.

**Fig 2.**
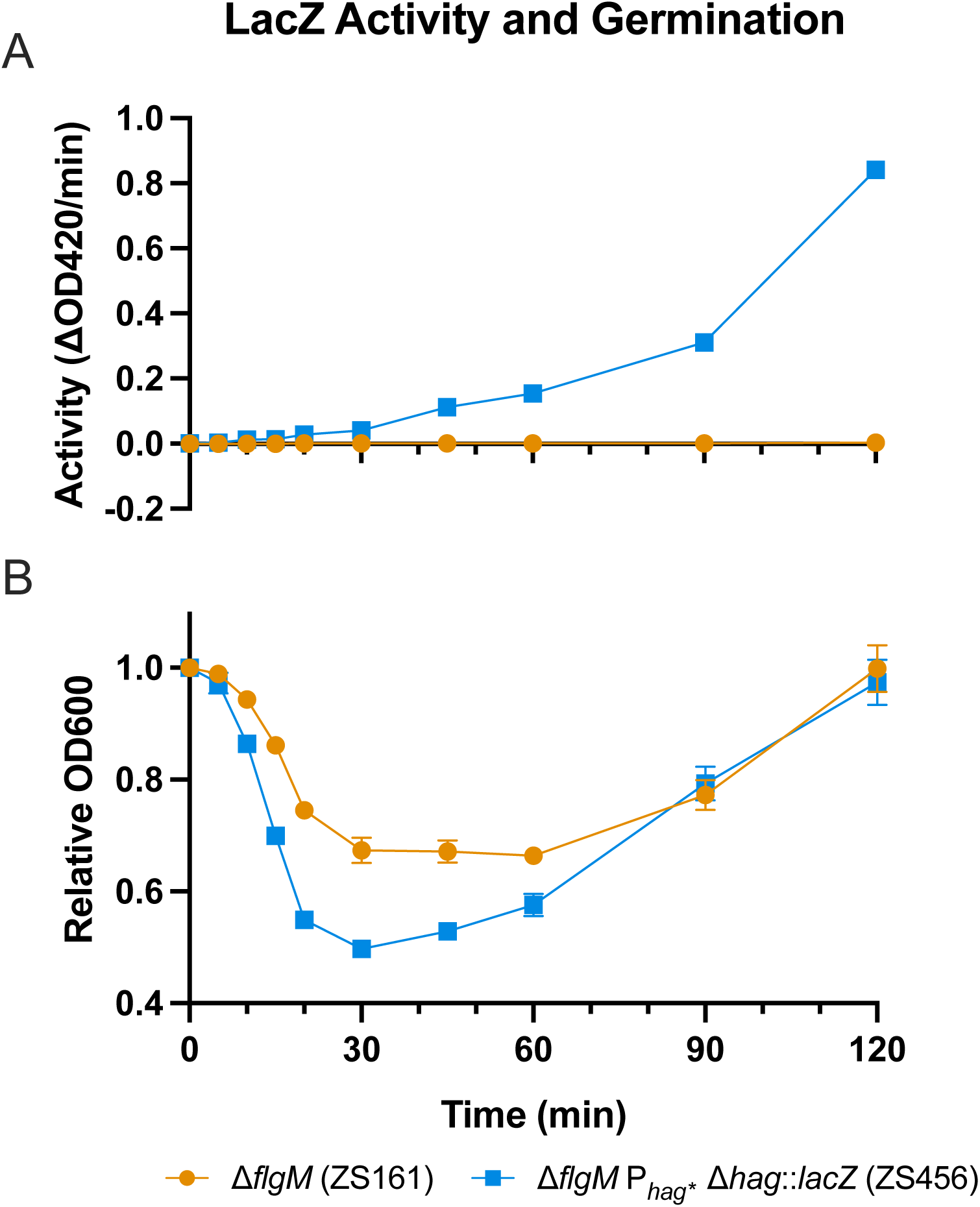
Engineered promoter enables robust protein expression. The efficiency of the P*_hag*_* promoter was assessed by quantification of expression in a LacZ reporter strain. **A**) LacZ activity of ZS161 (orange circles) and ZS456 (blue squares) was determined at each time point by measuring the change in absorbance at 420 nm following the conversion of ONPG to ONP. Light scattering corrections were made using the formula: Abs420-(1.75*Abs550). Activity was calculated as the rate over the linear range. **B**) Germination of ZS161 (orange circles) and ZS456 (blue squares) was evaluated by the change in OD_600_ and normalized using the OD_600_ obtained at time zero [relative OD_600_ = OD_600_(t)/OD_600_(t_0_)]. The data represent the averages from three independent measurements, and error bars represent the standard deviations (SD). If bars are not visible the SD is smaller than the icon size. Results are representative of 2 experiments

### ALDH activity is robust in the *hag* locus

ZS183 (Δ*flgM* P*_hag*_* Δ*hag*::*acoD*), was constructed to stimulate ALDH activity under the modified hag promoter (P*_hag*_*). We measured activity from germinating cultures in a rich medium (Fig 3A). Germination was completed by T_45_, as demonstrated by the dip in OD_600_, after which OD_600_ rose during outgrowth and subsequent vegetative growth (Fig 3A Bottom). ALDH activity was detected at T_105_ and continued to rise linearly between T_105_ and T_180_, which is indicative of constitutive expression at this stage of outgrowth (Fig 3A Top).

**Fig 3.**
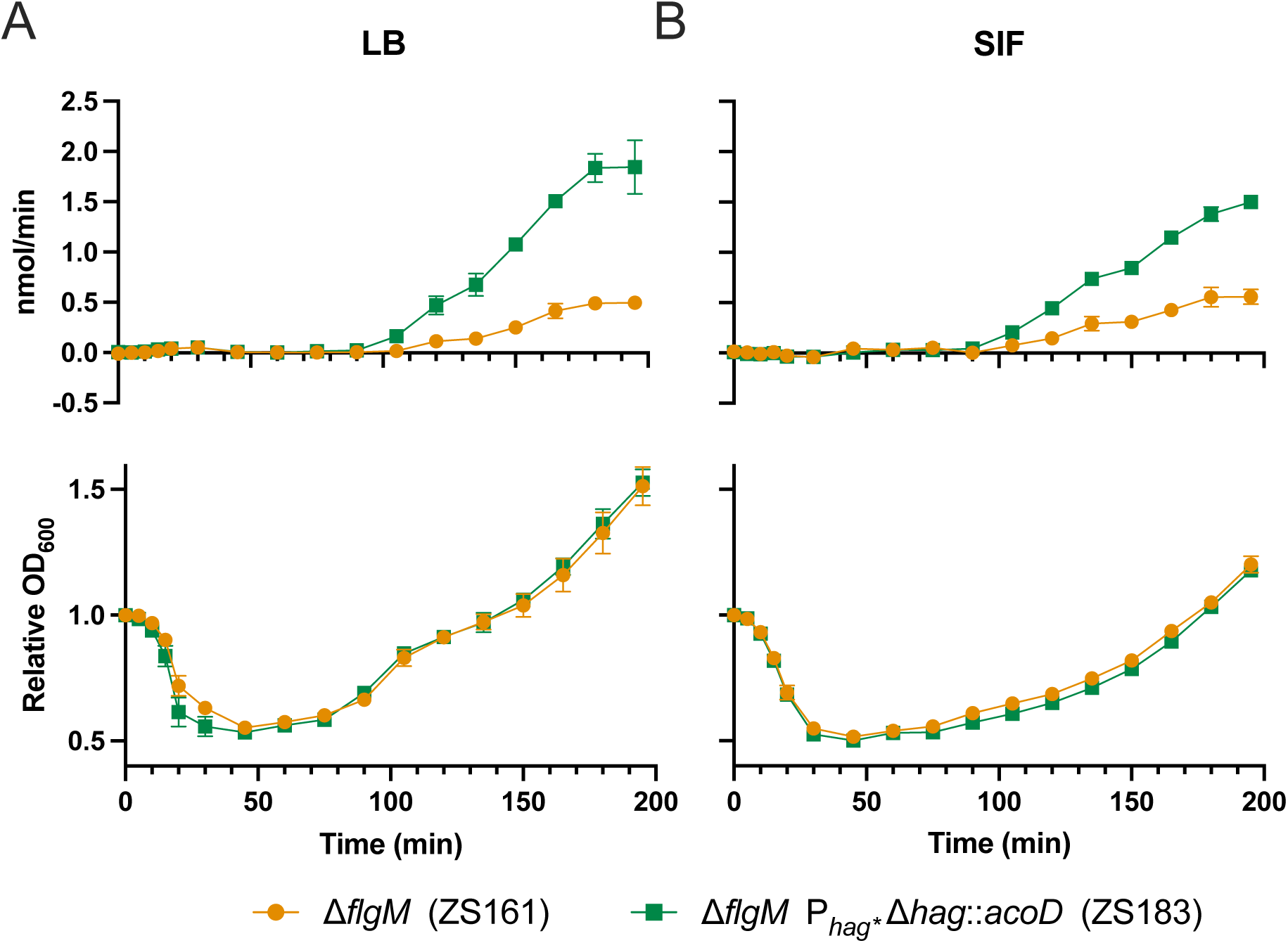
ALDH activity is robust in LB and SIF media. The expression of ALDH was evaluated under nutrient rich and nutrient limited conditions. **A) Top:** ALDH activity of ZS161 (orange circles) and ZS183 (green squares) in LB was determined by the change in absorbance at 340 nm following the production of NADH from NAD+ in the presence of AcA. The rate was calculated using the formula: nmol AcA/min = (ΔAbs340/εl)*v*1000. **Bottom:** Germination of ZS161 (orange circles) and ZS183 (green squares) in LB. **B) Top:** ALDH activity of ZS161 (orange circles) and ZS183 (green squares) in simulated intestinal fluid (SIF). **Bottom:** Germination of ZS161 (orange circles) and ZS183 (green squares) in SIF. The data represent the averages from 3 independent measurements, and error bars represent the standard deviations (SD). If bars are not visible the SD is smaller than the icon size. Results are representative of 2 experiments.

### ALDH activity is robust after germination in simulated intestinal fluid

We were interested in assessing whether conditions *in vivo* would impact germination rates and ALDH activity of engineered strains. To simulate conditions *in vivo*, we prepared a simulated small intestinal fluid (SIF) [41]. This media replicates the pH, salt stress, and bile acid stress of the small intestine where germination is likely to occur. Time to peak germination of spores in SIF was not significantly different from germination in rich media (T_45_), but outgrowth and cell division was slightly delayed likely due to the less rich media (Fig 3B Bottom). ALDH expression was detected at T_105_ about 1 hour after germination, similar to that seen in rich media (Fig 3B Top). Activity continued to increase thereafter up to T_195_. The absolute rate of activity is lower than in rich media, which is in line with slower outgrowth of the bacteria in SIF.

### Acetaldehyde removal is robust in whole cells

In the previously described experiments, we measured ALDH activity indirectly via the conversion of the enzyme’s cofactor NAD+ to NADH. To measure AcA levels directly, we adapted a protocol by Ressmann et al., for high-throughput quantification of AcA removal [42]. In addition, unlike the previous assays that relied on cell lysis to quantify activity of the deregulated *hag* promoter, we carried out the colorimetric assay on vegetative and germinating whole cells. Cells were grown up in LB and vegetative cells were rinsed and added to M9 minimal medium and spiked with AcA to a final concentration of 1mM. We then monitored the change in AcA over time by measuring the change in absorbance at 380 nm in the presence of 2--amino benzamidoxime (ABAO). AcA initially dropped 106 µM and 131 µM after exposure to vegetative whole cells of the ALDH-expressing strain ZS183 and the control strain ZS161, respectively, compared to 6 µM in the media only control, which we attribute to initial AcA interaction with the cells. From T_3_-T_27_ we recorded a 271 µM drop in AcA from ZS183, which was 4-fold higher than the 68 µM drop observed from ZS161 by T_27_ and 8-fold higher by T_54_ (622 µM and 75 µM, respectively). Overall, ZS183 removed >99% of AcA compared to 16.5% for ZS161 (Fig 4A).

**Fig 4.**
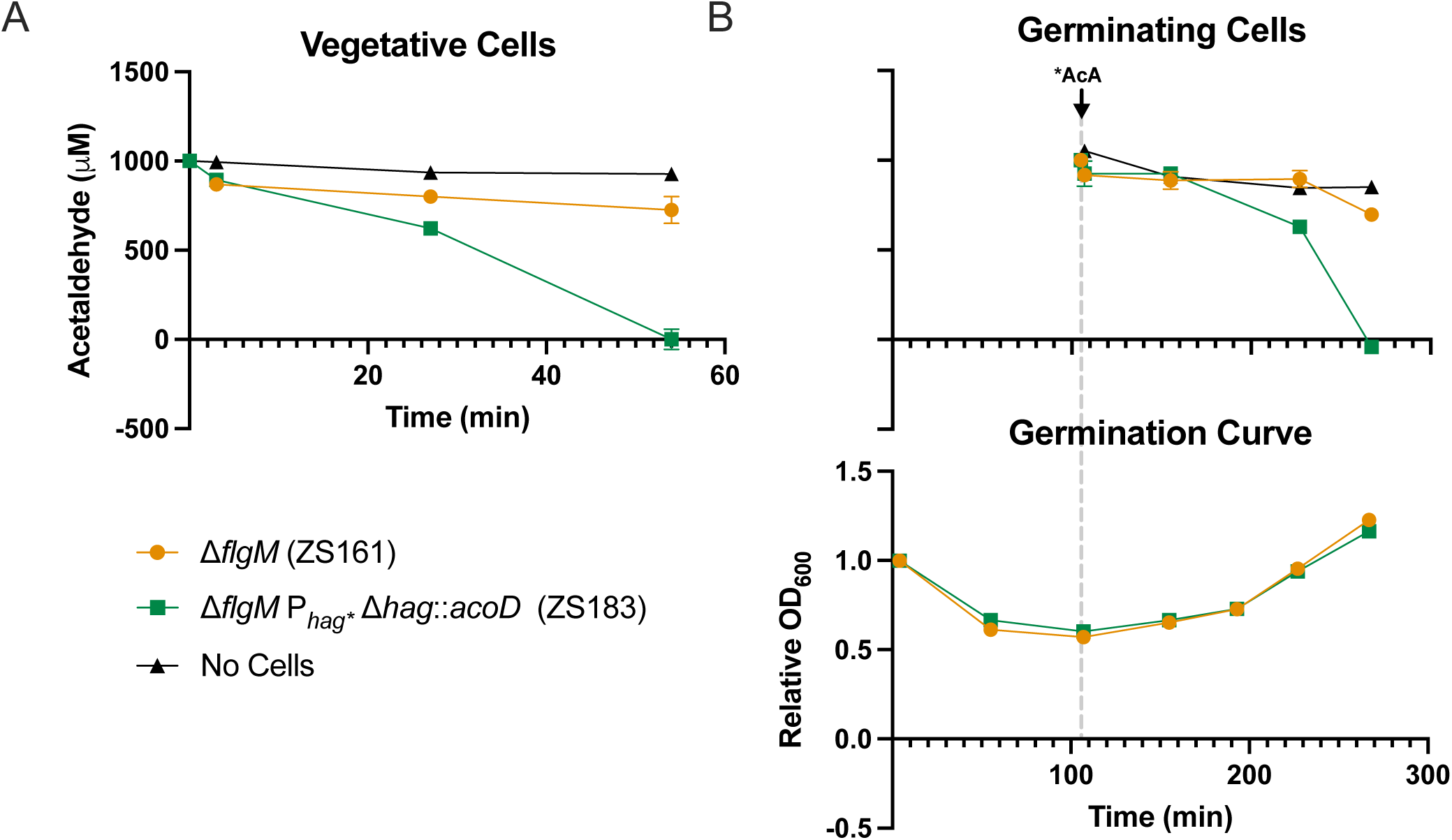
Engineered strain ZS183 efficiently removes acetaldehyde. The removal of acetaldehyde (AcA) by whole cells was quantified by the change in absorbance at 380 nm in the presence of ABAO. **A)** AcA removal by vegetative cells of ZS161 (orange circles) and ZS183 (green squares). **B) Top:** AcA removal by newly germinated, outgrowing cells of ZS161 (orange circles) and ZS183 (green squares). **Bottom:** Germination of ZS161 (orange circles) and ZS183 (green squares). Arrow and dotted line indicate when AcA was spiked into the germinating culture. Data represents the average of three independent measurements, and error bars represent the standard deviations (SD). If bars are not visible the SD is smaller than the icon size.

### Acetaldehyde removal is active and robust after germination from spores

To determine if intracellular ALDH activity and NADH turnover are robust in a physiologically relevant timeframe for a probiotic after germination, the whole-cell AcA removal assay using ABAO was repeated following germination of spores in rich media, rather than starting directly with vegetative cells. Spore germination was completed by T_60_ for both strains (Fig 4B Bottom). Due to the reactive nature of AcA we opted to spike it into the germinating population at T_105,_ just before the cells reached the stage of outgrowth where we previously detected *hag* promoter activity in the LacZ and ALDH time courses (Figs 2 and 3). The concentration of AcA remained unchanged in both cultures up to T_150_, as expected given the expression timeline of σ^D^ dependent genes during germination and outgrowth. By T_220,_ ALDH-expressing strain ZS183 removed approximately 14-fold more AcA than the control strain ZS161 (296 µM and 21 µM, respectively) and by T_270,_ approximately 4-fold more AcA than the control (925 µM and 219µM, respectively). Overall, it took 165 minutes for ZS183 to completely deplete the concentration of AcA added (Fig 4B Top). The rate of reduction of acetaldehyde was ∼41-fold faster in ZS183 than ZS161, 4.52 nmol/ml/min to 0.11 nmol/ml/min, respectively.

### Vegetative growth and germination are unaffected by heterologous protein expression in the constitutive hag promoter

For enzyme delivery *in vivo*, it is important that vegetative cell growth and spore germination is not impacted by: the burden of constitutive expression at the *hag* locus, changes in regulation from increased σ^D^ activity due to the *flgM* knockout, or effects of the expressed enzyme. While we demonstrated activity and germination in the previous experiments, we wanted to screen all the strains simultaneously to confirm there were no defects in growth or germination in engineered strains as compared to the parent strain of *B. subtilis*, PY79. Indeed, we did not observe any significant changes in growth (lag phase, exponential growth, stationary phase times) or germination (phase bright to phase dark germination, ripening, outgrowth) (Fig S1).

## Discussion

In this study we demonstrated a new constitutive expression system in relevant conditions that relies on derepression of the *hag* locus and its endogenous promoter for high levels of heterologous expression in place of the *hag* gene. Using this system, we engineered a first of its kind acetaldehyde (AcA) degrading probiotic capable of being ingested as a spore. Translational use-cases of a spore-forming probiotic can leverage mature endospore stability for oral delivery, storage convenience, and extended shelf life. With that in mind, we measured activity of the derepressed *hag* promoter during germination and outgrowth to confirm that constitutive expression of the *hag* locus was robust despite first undergoing sporulation. Through the repeat detection of increasing activity over time, we demonstrated that expression is robust after germination. This is especially crucial as it relates to the translational application of a probiotic that is delivered as a spore, where expression needs to be robust shortly after germination in the intestinal microenvironment.

We relied on absorbance at 600 nm to track the germination timeline. At the timepoints we assessed, changes in OD_600_ are not indicative of increases or decreases in total cell count, but rather they mark the physiological stages of germination and outgrowth [43]. As such, we did not normalize the data based on absorbance as a proxy for overall population density. To ensure activity was based on heterologous protein expression and not population density, we started the assays with the same spore OD_600_ for each sample.

In our first assessment of the derepressed *hag* promoter, we built a LacZ reporter strain (ZS456) and detected activity at T_45_ post germination-initiation, which is in line with the expression timelines of early activated cellular processes previously observed during *B. subtilis* spore revival (Fig 2A) [40]. In our engineered AcoD-expression strain (ZS183) we detected initial activity at T_105_, close to an hour later than that observed in the LacZ expression strain (Fig 3). This was likely due to a higher limit of detection in the ALDH assay and matches up with the rise in LacZ activity measured at T_90_-T_120._

Survival of the probiotic chassis and robust *in vivo* germination are essential for achieving enzymatic function in the body. Previous scientific research demonstrates the capacity for *B. subtilis* spores to germinate continuously over the course of 24 hours in the small intestines of ileostomy patients, without adverse impacts [17]. To simulate strain activity in the gastrointestinal tract, we demonstrated ALDH activity from recently germinated spores in simulated small intestinal fluid (SIF). Spores cultured in both LB and SIF showed robust constitutive expression upon maturation of germinated spores. However, since germination and outgrowth are optimally carried out in rich media, we detected earlier outgrowth compared to the same assay in SIF media (Fig 3).

The ALDH activity experiments in figure 3 were done using cell lysates, and so we wanted to ensure that the same results could be seen in whole cells, especially without the addition of the necessary cofactor NAD+. The Δ*flgM* control strain without AcoD showed that *B. subtilis* has minimal endogenous AcA dehydrogenase activity characterized by a slight reduction in AcA as compared to wells without whole cells and is likely due to non-specific activity of a general aldehyde dehydrogenase or other enzyme(s) with related function (Fig 4). In both vegetative and germinating AcoD expressing cells, AcA concentrations rapidly fall below a detectable limit, further confirming that the expression of ALDH under P*_hag_*_*_ is robust and capable as a probiotic designed for gut-derived AcA removal.

*B. subtilis* is an attractive candidate for the probiotic space due to the species’ safety, stability, and ease of engineering. However, limitations in heterologous protein expression based on genomic integration versus high copy number plasmids as well as protease expression have lessened the popularity of this organism in the synthetic biology space. For applications where steady expression and stability are favored, *B. subtilis* has an edge on common engineering approaches such as traditional lysis circuits and its ability to sporulate provides natural protection and encapsulation. Here we are adding to the engineering toolbox by presenting a novel site of integration with an endogenous promoter and showing its capacity to translate into real world use cases.

Since this genetically engineered strain of *B. subtilis* is targeted for human consumption and therefore exposed to a complex intestinal microbiome, we intentionally designed our chassis to minimize the potential for interactions within the gut microenvironment. Introduction of the *acoD* gene from *C. necator* is the single addition of exogenous DNA in ZS183 (Δ*flgM* P*_hag*_* Δ*hag*::*acoD*). This gene was specifically selected from a strain that has historically colocalized with *B. subtilis* in soil environments. Given the long timeline (hundreds of millions of years) that these two species have been in contact, and the sheer number of cells that have interacted over that timeline, it is far more likely than not that *acoD* has been acquired by *B. subtilis* and subsequently lost over time, probably many times over. Importantly the function of the gene is identical to the function of other endogenous genes in *B. subtilis* [44]. Although highly similar to known food-safe genes, the expressed product encoded by *C. necator’s acoD* does not specifically have a well-established history in human food. To alleviate any safety concerns, we piloted a 90-day repeated oral toxicity in rats that showed no adverse effects on clinical health [45]. The engineered strain was sequenced and shown to be free of virulence factors/toxins associated with pathogenicity and of transferable antibiotic resistance genes, which matches the known traits of *B. subtilis* PY79. We also had tests conducted for hemolysis, antibiotic resistance (minimum inhibitory concentration), and cell cytotoxicity with no adverse observations.

The engineered strain ZS183 has now been brought to market (ZB183^TM^) – transparently labeled as “proudly GMO” – to positive public reception. Public adoption of an engineered probiotic that delivers a benefit to consumers directly has been strong, with over 4 million bottles sold as of the date of this writing. This is consistent with a study of consumer GMO sentiment, which found that the majority of consumers polled believed that others were ‘against GMOs’ while they themselves were not opposed to a product that benefitted them directly [46]. This implies that there is a rich future of improvement and benefit available through the rational engineering of safe and active probiotics such as that described in this study.

## Materials and Methods

### Bacterial strain and growth conditions

All engineered strains described in this work are derived from *Bacillus subtilis* PY79. *B. subtilis* strains were routinely cultured at 37°C with shaking at 250 rpm or grown on agar plates at 37°C in aerobic environments. Stellar *E. coli* HST08 was used for subcloning. *E. coli* was cultured at 37°C with shaking at 250 rpm or grown on agar plates at 37°C in aerobic environments. Media was supplemented with ampicillin (100 µg/ml) and MLS (25 µg/ml lincomycin and 1 µg/ml erythromycin) when necessary.

### Plasmid construction

To enable expression at the *hag* locus, homologous regions of *hag* were cloned into pMiniMAD which has a temperature-sensitive origin of replication and ampicillin and mls resistance markers for selection in *E. coli* and *B. subtilis*, respectively [47]. All plasmids and primers used in this study are listed in Tables 1 and 2, respectively. Briefly, a gBlock (ZP72) containing *GFPmut3* (*gfp*) flanked by 800 bp of upstream and downstream *hag* sequence homology was synthesized by Integrated DNA Technologies (IDT). The 5’ homology region carries a point mutation (G → A) in the *hag* promoter at position −38 relative to the start codon of *hag* to disrupt binding of CsrA (P*_hag*_*). The gBlock, ZP72 was cloned into pMiniMAD linearized by primers ZP24F and ZP25R to generate pZB149 (P*_hag*_ Δhag*::*gfp*). All subsequent *hag* locus integration vectors were constructed from p149 by replacing GFP with the gene of interest (GOI). To generate pZB163 (P*_hag*_ Δhag*::*acoD*), *acoD* codon optimized to *B. subtilis* and synthesized by Genscript was amplified by primers ZP83F and ZP82R and cloned into pZB149 linearized by PCR using primers ZP80F and ZP79R. To generate pZB413 (P*_hag*_ Δhag*::*lacZ*) *lacZ* was amplified using primers ZP203F and ZP204R and cloned into pZB149 linearized by PCR using primers ZP88F and ZP91R. Homologous regions 800 bp upstream and downstream of *flgM* were synthesized by IDT and cloned into pMiniMAD linearized using primers ZP24F and ZP25R to generate pZB147 (Δ*flgM*). All plasmids were confirmed by PCR and Sanger sequencing using primers outlined in Table 2.

**Table 1.** Bacterial strains and plasmids used in this study.

| Bacterial Strain or Plasmid | Relevant Background | Source/Reference |
| --- | --- | --- |
| **Plasmids** |  |  |
| pMiniMAD2 | oriBsTs amp<sup>r</sup>, er<sup>r</sup>, mls<sup>r</sup> | Parashar et al., 2013; Patrick et al, 2008 |
| pZB147 | $\Delta flgM$ | This study |
| pZB149 | $P_{hag^*} \Delta hag::gfp$ | This study |
| pZB163 | $P_{hag^*} \Delta hag::acoD$ | This study |
| pZB413 | $P_{hag^*} \Delta hag::lacZ$ | This study |
| **B. subtilis** |  |  |
| PY79 (ZS6) | *swrA sfpO* | Youngman, Perkins, & Losick, 1984 |
| ZS161 | $\Delta flgM$ | This Study |
| ZS180 | $\Delta flgM P_{hag}^*$<br>$\Delta hag::gfp$ | This Study |
| ZS183 | $\Delta flgM P_{hag}^*$<br>$\Delta hag::acoD$ | This Study |
| ZS456 | $\Delta flgM P_{hag}^*$<br>$\Delta hag::lacZ$ | This Study |

**Table 2.**
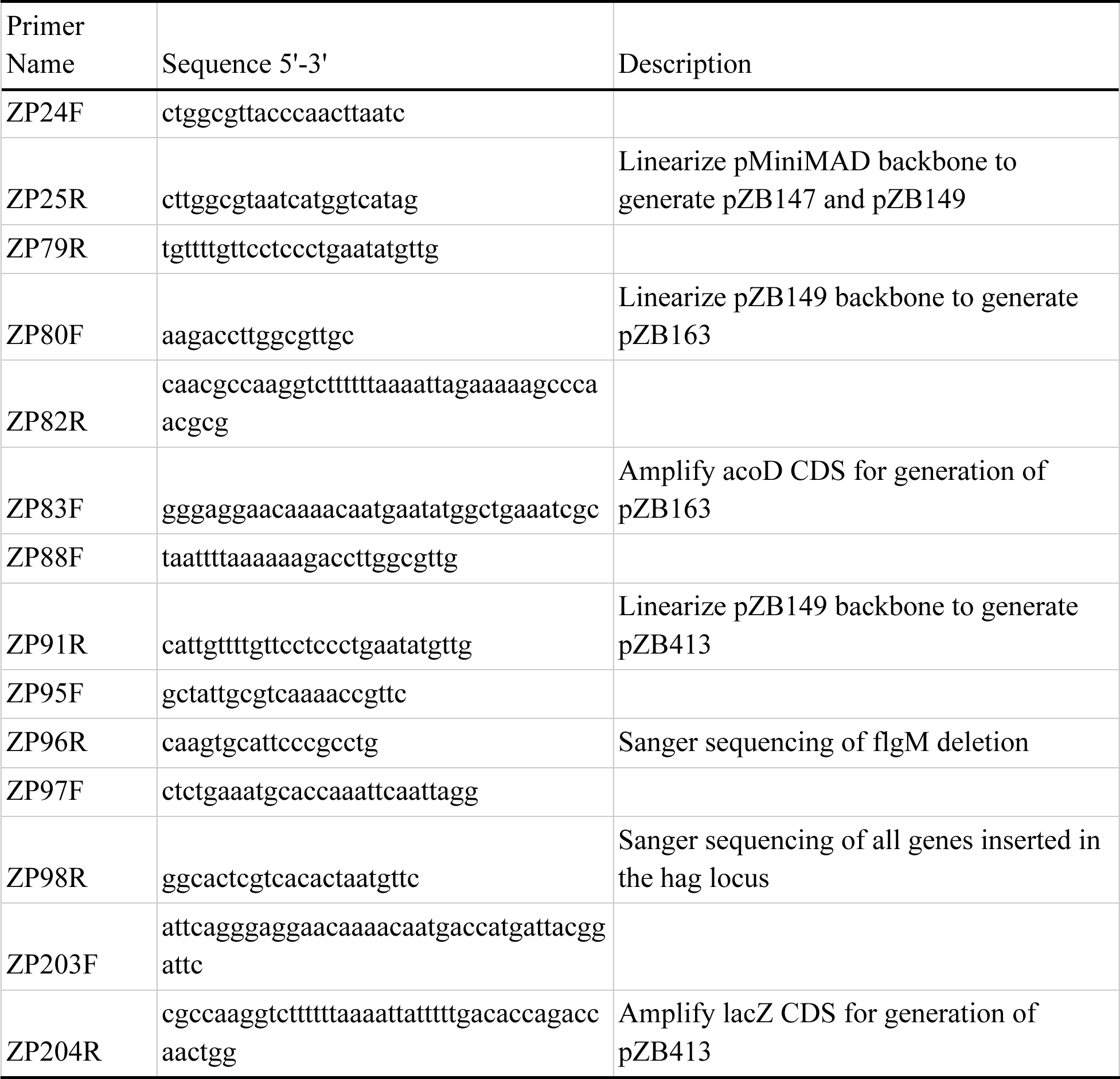
Primers used in the present study.

### Engineered *B. subtilis* strain construction

Engineered strains of *B. subtilis* were generated by natural competence. Briefly, strains of *B. subtilis* were streaked out on LB agar to yield single colonies. Liquid cultures were prepared by inoculating modified competence (MC) media (0.1 M K_3_PO_4_ pH 7.0, 3 mM Na_3_C_6_H_5_O_7_, 3 mM MgSO_4_, 22 mg/mL ferric ammonium citrate, 2% glucose 0.1% casein hydrolysate, 0.2% potassium glutamate) with a single colony and growing to OD_600_ >1.1. From this culture, 400 µl of cells were mixed with >1 µg plasmid DNA and incubated at the restrictive temperature (37°C) with shaking for approximately 2 hours. Cultures were then plated on agar supplemented with MLS (25 µg/ml lincomycin and 1 µg/ml erythromycin) and incubated overnight at 37°C. Colonies that grew overnight were assumed to be merodiploids. To stimulate recombination and resolve the merodiploid due to the temperature-sensitive origin of replication in the pMiniMad plasmid, liquid cultures were prepared by inoculating LB broth with a single colony and growing overnight at the permissive temperature (25°C) with shaking. Overnight cultures were subcultured in fresh LB broth and incubated for 8-12 hours at 25°C with shaking three times. The final subculture was plated on LB agar and incubated overnight at 37°C. Single colonies were replica patched on LB agar and LB agar supplemented with MLS. MLS sensitive cured recombinants were PCR-screened for integration and confirmed by Sanger sequencing.

*B. subtilis* PY79 was transformed with pZB147 and cured to yield ZS161 (Δ*flgM*). Subsequently ZS161 was transformed with plasmid pZB149 and cured to yield strain ZS180 (Δ*flgM* P*_hag*_Δhag*::*gfp*). To create the AcoD-expressing strain, ZS161 was transformed and cured of pZB163 to generate ZS183 (Δ*flgm* P*_hag*_* Δ*hag*::*acoD*). The LacZ reporter strain was constructed by transformation and curing of ZS161 with pZB413 to generate ZS456 (Δ*flgM* P*_hag*_* Δ*hag*::*lacZ*).

### Analysis of vegetative cell growth

Vegetative cell growth was monitored by the change in absorbance at 600 nm (optical density, OD_600_) using a Synergy H1 microplate reader (BioTek; Gen5 v3.10 software). Frozen stocks of *B. subtilis* strains were streaked onto LB agar (Lennox; RPI) plates and incubated overnight at 37°C to yield single colonies. Liquid cultures of *B. subtilis* strains were prepared by inoculating LB broth (Lennox; RPI) with a single colony and growing overnight at 37°C with shaking (250 rpm). The following day, strains were subcultured into fresh LB broth to an OD_600_ = 0.1. Aliquots (200 µl) of each culture were added to a 96-well plate, n=10. The OD_600_ was measured once every 20 minutes for 8 hours. After background subtraction, OD_600_ values were plotted to generate growth curves.

### Spore preparation and purification

Frozen stocks of *B. subtilis* strains were streaked onto nutrient agar (Difco) plates and incubated overnight at 37°C to yield single colonies. Liquid cultures of *B. subtilis* strains were prepared by inoculating nutrient broth (RPI) with a single colony and grown at 37°C with shaking (250 rpm) until mid-exponential phase of growth. Aliquots (200 μl) from the liquid *B. subtilis* cultures were spread onto sporulation media (nutrient agar supplemented with 10% KCl, 1.2% MgSO_4_, 1M Ca(NO_3_)_2_, 10mM MnCl_2_, and 1mM FeSO_4_) and incubated at 37°C for five days to allow for sporulation. The resulting bacterial lawns were harvested by scraping into ice-cold RO water. The spores were pelleted by centrifugation at 16,000xg for 1 minute. The supernatant was decanted, and the spore pellet was resuspended in RO water. This process was repeated three times.

Spores were purified by density centrifugation through a 20%-50% Histodenz gradient. Following the third wash with RO water, the spore pellet was resuspended in a 20% Histodenz solution, layered on top of the 50% Histodenz solution, and centrifuged at 21,000xg for 5 minutes. After discarding the Histodenz solution, the resulting spore pellet was washed three times by repeated centrifugation (16,000xg, 1 minute) and resuspension in RO water. Purity was assessed by phase contrast microscopy for the appearance of phase bright spores and the absence of vegetative cells. The spores were stored in sterile RO water at 4°C until needed.

Where indicated, spores were alternatively prepared using a chilled-plate method. In brief, 100 µl of mid log culture was plated on LB agar plates and grown at 37°C for 72 hours to allow for complete sporulation. Plates were then placed at 4°C for 4-7 days to allow for natural autolysis by *B. subtilis* vegetative cells. Spores were purified from cellular debris by washing in sterile RO water and pelleting by centrifugation at 16,000xg for 2 minutes for at least 3 washes. Purity was confirmed by microscopy for absence of vegetative cells.

### Analysis of spore germination

Spore germination was monitored by the decrease in absorbance at 600 nm (optical density, OD_600_) using an Epoch microplate reader (BioTek, Gen5 v3.10 software). Aliquots of purified spores were suspended in sterile RO water to OD_600_ = 1.0 and concentrated 10X. Germination assays were carried out in 96-well plates at a 360 µl/well final volume. Germination reactions were performed in triplicate wells and consisted of 324 µl LB supplemented with 10 mM alanine and 36 µl of the 10X spore suspension. The initial OD_600_ of the reaction was ∼1.0. The OD_600_ was measured once every minute for two hours and normalized using the OD_600_ obtained at time zero [relative OD_600_ = OD_600_(t)/OD_600_(t_0_)]. Resulting curves were analyzed for three metrics: peak rate of percent change in OD_600_ (r_max), time to reach peak rate (t_max), and overall percent drop in OD_600_ (%dOD). An up to 60% drop in OD_600_ can be interpreted as effective germination [43].

### Analysis of LacZ activity

LacZ expression during spore outgrowth was measured in cell lysates by the change in absorbance at 420 nm following the conversion of ONPG to ONP using an Epoch microplate reader (BioTek; Gen5 v3.10 software). Reagents were prepared as follows: assay buffer (0.06 M Na_2_HPO_4_⋅7H_2_O, 0.04 M NaH_2_PO_4_⋅H_2_O, 10 mM KCl, 1 mM MgSO_4_, pH 7.0); BugBuster Lysozyme (BBL) solution (99% BugBuster Protein Extraction reagent (Millipore) and 1% egg white lysozyme (GoldBio) solution (20 mg/ml in 50 mM Tris, pH 8.8)); lysis buffer (90% assay buffer and 10% BBL); ONPG solution (4 mg/ml).

Aliquots of purified spores were suspended in sterile RO water to OD_600_ = 1.0 and concentrated 10X. Flasks containing LB broth were inoculated with 10X spore suspensions to OD_600_ 1.2 and incubated at 37°C with shaking (250 rpm). At time zero (T_0_), 1 mL of the culture was removed and transferred to a chilled microtube on ice. From this sample, 180 µl was transferred to a microplate well containing 180 µl LB, mixed by pipetting, and its D_600_ measured. The remaining cells were pelleted by centrifugation at 16,000xg for 1 minute, the media was decanted, and the pellet stored at −80°C until needed. These steps were repeated at each time point.

Frozen pellets were resuspended 1:1 in the lysis buffer and incubated at room temperature for 10 minutes. LacZ activity was measured in 96-well plates. Reactions consisted of 60 µl of assay buffer and 100 µl of cell suspension. Absorbance was first measured at 420 nm and 550 nm. Next, 50 µl of the ONPG solution was added to each well. Absorbance readings at 420 nm and 550 nm were then taken every minute for 10 minutes. Light scattering at 420 nm due to cell debris was corrected using the formula: Abs420-(1.75*Abs550). Activity was then calculated as the rate over the linear range with final units of ΔAbs420/min.

### Analysis of ALDH expression in complex media

ALDH expression during spore outgrowth in complex media was measured in cell lysates by the change in absorbance at 340 nm following the production of NADH from NAD+ in the presence of AcA using an Epoch microplate reader (BioTek; Gen5 v3.10 software). Reagents were prepared as follows: 50 mM Tris buffer (pH 8.8); 100 mM NAD+ (in 50 mM Tris buffer, pH 8.8); 50 mM AcA (in RO water); BugBuster Lysozyme (BBL) solution (99% BugBuster Protein Extraction reagent (Millipore) and 1% egg white lysozyme (GoldBio) solution (20 mg/ml in 50 mM Tris, pH 8.8)).

Aliquots of purified spores were suspended in sterile RO water to OD_600_ = 1.0 and concentrated 10X. Flasks containing LB broth were inoculated with 10X spore suspensions to OD_600_ 1.2 and placed in a shaking incubator (250 rpm) at 37°C. At time zero (T_0_), 1 mL of the culture was removed and transferred to a chilled microtube on ice. From this sample, 180 µl was transferred to a microplate well containing 180 µl LB, mixed by pipetting, and its OD_600_ measured and normalized using the OD_600_ obtained at time zero [relative OD_600_ = OD_600_(t)/OD_600_(t_0_)]. The remaining cells were pelleted by centrifugation at 16,000xg for 1 minute, the media was decanted, and the pellet stored at −80°C until needed. These steps were repeated at each time point.

Frozen pellets were resuspended in 200 µl BBL and incubated at room temperature for 10 minutes. Cells were pelleted by centrifugation at 16,000xg for one minute and the lysate transferred to a chilled microtube on ice. Assays were carried out in 96-well plates at a 360 µl/well final volume. Each well was prepared with 264 µl Tris buffer, 36 µl NAD+, and 50 µl cell lysate. Reactions were initiated by the addition of 10 µl AcA. Absorbance at 340 nm was measured every minute for 10 minutes. The peak rate averaged over 4 minutes was then converted to nmol/min AcA using the following equation and a 1:1 ratio of NADH to AcA 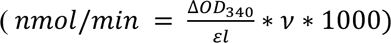 where ΔOD_340_ is the change in OD_340_ per minute, *ɛ* is the extinction coefficient for NADH (6220 M^-1^cm^-1^), *l* is the path length (1 cm), and *v* is the reaction volume (360 µl).

### Analysis of ALDH activity in simulated intestinal media

ALDH expression during spore outgrowth in simulated intestinal media (16.5 g/L tryptone (RPI) supplemented with FaSSIF Buffer Concentrate (Biorelevant; 3 mM sodium taurocholate, 0.75 mM phospholipids, 148 mM sodium, 106 mM chloride, 29 mM phosphate, pH 6.5)) was measured in cell lysates as described above for ALDH activity in complex media.

### Analysis of acetaldehyde removal by vegetative cells

Acetaldehyde (AcA) removal by vegetative cells was measured by sampling supernatant and assaying AcA using the change in absorbance at 380 nm in the presence of 2--amino benzamidoxime (ABAO), (as inspired by aldehyde quantifications by Ressmann et. al, 2019) using an Epoch microplate reader (BioTek; Gen5 v3.10 software) [42]. Reagents were prepared as follows: M9 minimal media (0.24 M Na_2_HPO_4_•7H_2_O; 0.11 M KH_2_PO_4_; 0.043 M NaCl; 0.093 M NH_4_Cl); sodium acetate buffer (100 mM, pH 4.5); ABAO solution (2.5 mM in sodium acetate buffer)

Frozen stocks of *B. subtilis* strains were streaked onto LB agar plates and incubated overnight at 37°C to yield single colonies. Liquid cultures of *B. subtilis* strains were prepared by inoculating LB broth with a single colony and growing at 37°C with shaking for 6.5 hours. The cells were pelleted by centrifugation at 3,200xg for 10 minutes, the media was decanted, and the pellet washed twice with M9 media. Next, cells were suspended in M9 media to OD_600_ = 1.400 ± 0.050. The change of media was necessary to decrease background interference.

To further account for background interference, each sample was split into two cultures of equal volume. AcA was added to one culture at final concentration of 1 mM, sterile RO water was added to the other culture. Cultures were then placed in an incubator at 30°C with shaking at 250 rpm. At each time point, an aliquot (200 µl) was removed from each culture and pelleted by centrifugation at 16,000xg for 2 minutes. The supernatant was then transferred to a chilled microtube on ice and immediately quantified in an ABAO assay.

ABAO assays were carried out in 96-well plates at a 350 µl/well final volume. Reactions consisted of 150 µL sodium acetate buffer and 150 µl sample supernatant. A standard curve from 1 mM to 62.5 µM of AcA in M9 media was included on each plate. An initial absorbance measurement at 380 nm was taken. Next, 50 µl of ABAO solution was added to each well. Subsequently, the absorbance at 380 nm was measured once every minute for 20 minutes. In all cases the value at T_20_ was used as final absorbance values for calculations.

Concentrations of AcA were calculated by first subtracting the initial absorbance (380 nm in all cases) of each sample from the final absorbance values for each sample. Next, the absorbance of samples without AcA was subtracted from the absorbance of samples with AcA. For the standard curve a media only well was subtracted from the rest of the curve. The remaining absorbance is attributed to the level of AcA in the sample and the concentration was determined using the internal standard curve. A small background reaction wherein AcA reduces ABAO background reactions was unable to be accounted for and causes some results to drop slightly below zero (the control in which AcA was not added has higher background absorbance than the identical reaction in which AcA has been added and then enzymatically removed). For transparency we have chosen to present these negative values ‘as is’.

### Analysis of acetaldehyde removal by outgrowing cells

AcA removal during spore outgrowth was measured by sampling supernatant and assaying AcA using the change in absorbance at 380 nm in the presence of ABAO using an Epoch microplate reader (BioTek; Gen5 v3.10 software). Liquid cultures of each sample were prepared by inoculating LB broth (supplemented with 10 mM alanine) with spores prepared by the chilled plate method to OD_600_ = 1.1 ± 0.05. Each sample was then split into two cultures of equal volume and incubated at 37°C with shaking (250 rpm). At T_105_ AcA was added to one culture to a final concentration of 1mM; an equal volume of sterile RO water was added to the other culture. This was repeated for all samples. At each time point, an aliquot was removed from each culture. OD_600_ was recorded and the remaining sample volume was pelleted by centrifugation at 16,000xg for 2 minutes. ABAO assays were performed on the supernatant and AcA concentrations calculated as described above. To calculate rates, the amount of AcA removed between T_107_ and T_267_ was divided by the time between those two points (150 minutes). To account for the effect of evaporation, the rate of AcA lost in the media-only control over the same time period was subtracted from the rates in samples containing cells. The remaining rates were used to quantify the effects of the cells in each sample.

## Acknowledgements

We would like to express our sincere gratitude to Professor Dan Kearns from Indiana University Bloomington for generously providing the PY79 strain and pMiniMAD2 plasmid used in this study. We are also deeply appreciative of their insightful discussions throughout the course of this research.

## Notes

### Competing Interest Statement

CH, VR, BS, JO, and ZA are employees of ZBiotics Company and may hold stock/stock options in ZBiotics Company

