## Supporting Information for "Engineering a probiotic *Bacillus subtilis* for acetaldehyde removal: A *hag* locus integration to robustly express acetaldehyde dehydrogenase"

**S1 Fig. Growth and germination of strains used in this study.** The germination and growth profiles of the parent and engineered strains were evaluated by changes in $OD_{600}$. **A)** Growth curve of the wildtype parent strain PY79 (black circles) compared to the engineered strains. The data represent the averages from at least three independent measurements, and error bars represent the standard deviations (SD). If bars are not visible the SD is smaller than the icon size. **B)** Germination of PY79 (black circles) compared to the engineered strains. The data represent the averages from at least three independent measurements, and shaded areas represent SD.

**S1 Table. Germination metrics in all strains.**

**S2 Fig. Germination of strains in SIF media.** Germination at room temperature of ZS161 (orange line) and ZS183 (green line) in SIF was evaluated by changes in $OD_{600}$ of 360 µl cultures, using an Epoch microplate reader (BioTek; Gen5 v3.10 software). The data represent the averages from three independent measurements, and the shaded area represents the standard deviations (SD).

**S2 Table. Germination metrics in SIF vs LB.**

**S1 Fig. Growth and germination of strains used in this study.**

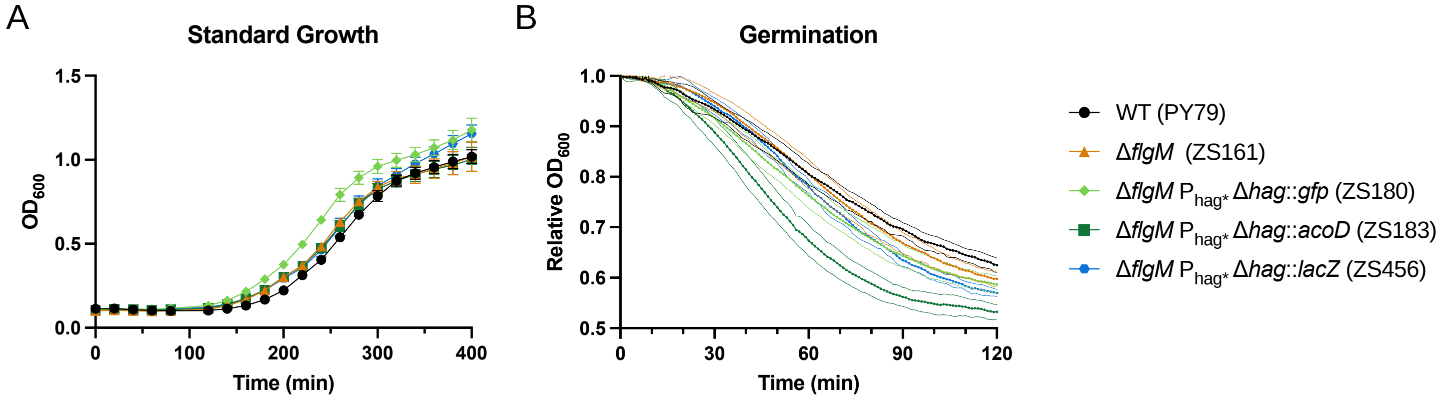


**S1 Table. Germination metrics in all strains.**

| | %dOD | t_max | %_max |
|---|---|---|---|
| **ZS6** | 36.6 ± 1.12 | 54.0 ± 5.66 | 0.542 ± 0.0589 |
| **ZS161** | 39.8 ± 0.957 | 59.7 ± 2.87 | 0.665 ± 0.0422 |
| **ZS180** | 40.8 ± 1.31 | 48.7 ± 4.99 | 0.629 ± 0.0573 |
| **ZS183** | 46.3 ± 1.17 | 41.3 ± 3.3 | 0.976 ± 0.0451 |
| **ZS456** | 42.5 ± 0.787 | 52.7 ± 1.89 | 0.762 ± 0.0105 |

**S2 Fig. Germination of strains in SIF media**

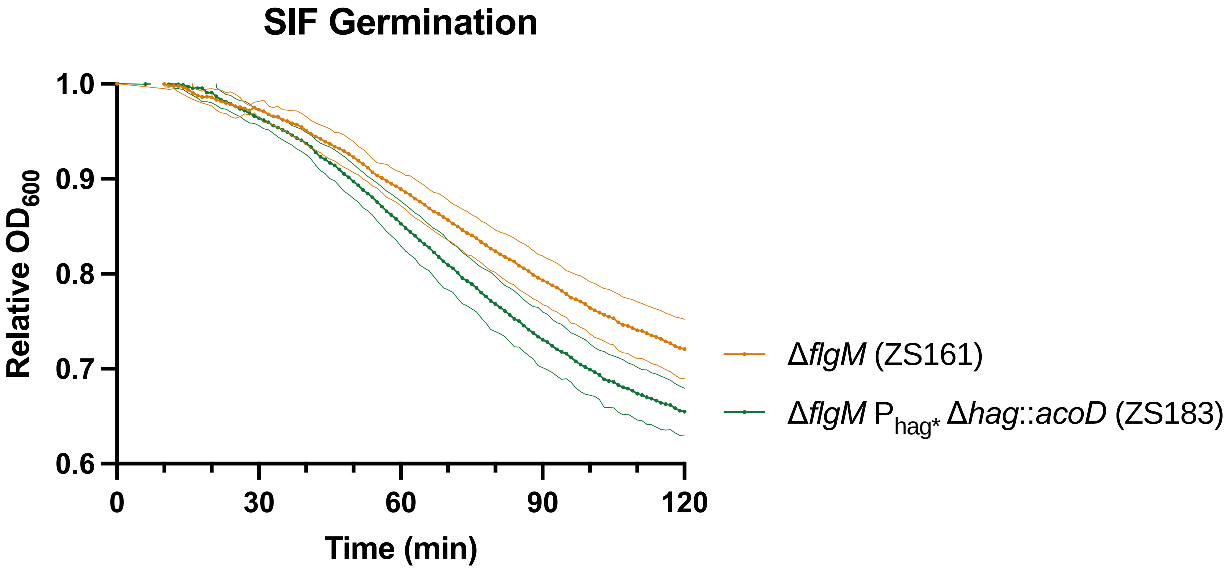


**S2 Table. Germination metrics in SIF vs LB.**

| | %dOD | t_max | %_max |
|---|---|---|---|
| **ZS161 LB** | 30.2 ± 1.05 | 60.3 ± 10.8 | 60.3 ± 10.8 |
| **ZS 161 SIF** | 27.4 ± 2.43 | 63.3 ± 3.3 | 63.3 ± 3.3 |
| **ZS183 LB** | 36.4 ± 2.2 | 62.3 ± 5.91 | 62.3 ± 5.91 |
| **ZS 183 SIF** | 34.0 ± 2.2 | 60.0 ± 3.56 | 60.0 ± 3.56 |
